# Thiopurines activate an antiviral unfolded protein response that blocks viral glycoprotein accumulation in cell culture infection model

**DOI:** 10.1101/2020.09.30.319863

**Authors:** Patrick Slaine, Mariel Kleer, Brett Duguay, Eric S. Pringle, Eileigh Kadijk, Shan Ying, Aruna D. Balgi, Michel Roberge, Craig McCormick, Denys A. Khaperskyy

## Abstract

Enveloped viruses, including influenza A viruses (IAVs) and coronaviruses (CoVs), utilize the host cell secretory pathway to synthesize viral glycoproteins and direct them to sites of assembly. Using an image-based high-content screen, we identified two thiopurines, 6-thioguanine (6-TG) and 6-thioguanosine (6-TGo), that selectively disrupted the processing and accumulation of IAV glycoproteins hemagglutinin (HA) and neuraminidase (NA). Selective disruption of IAV glycoprotein processing and accumulation by 6-TG and 6-TGo correlated with unfolded protein response (UPR) activation and HA accumulation could be partially restored by the chemical chaperone 4-phenylbutyrate (4PBA). Chemical inhibition of the integrated stress response (ISR) restored accumulation of NA monomers in the presence of 6-TG or 6-TGo, but did not restore NA glycosylation or oligomerization. Thiopurines inhibited replication of the human coronavirus OC43 (HCoV-OC43), which also correlated with UPR/ISR activation and diminished accumulation of ORF1ab and nucleocapsid (N) mRNAs and N protein, which suggests broader disruption of coronavirus gene expression in ER-derived cytoplasmic compartments. The chemically similar thiopurine 6-mercaptopurine (6-MP) had little effect on the UPR and did not affect IAV or HCoV-OC43 replication. Consistent with reports on other CoV Spike (S) proteins, ectopic expression of SARS-CoV-2 S protein caused UPR activation. 6-TG treatment inhibited accumulation of full length S0 or furin-cleaved S2 fusion proteins, but spared the S1 ectodomain. DBeQ, which inhibits the p97 AAA-ATPase required for retrotranslocation of ubiquitinated misfolded proteins during ER-associated degradation (ERAD) restored accumulation of S0 and S2 proteins in the presence of 6-TG, suggesting that 6-TG induced UPR accelerates ERAD-mediated turnover of membrane-anchored S0 and S2 glycoproteins. Taken together, these data indicate that 6-TG and 6-TGo are effective host-targeted antivirals that trigger the UPR and disrupt accumulation of viral glycoproteins. Importantly, our data demonstrate for the first time the efficacy of these thiopurines in limiting IAV and HCoV-OC43 replication in cell culture models.

**IMPORTANCE:** Secreted and transmembrane proteins are synthesized in the endoplasmic reticulum (ER), where they are folded and modified prior to transport. During infection, many viruses burden the ER with the task of creating and processing viral glycoproteins that will ultimately be incorporated into viral envelopes. Some viruses refashion the ER into replication compartments where viral gene expression and genome replication take place. This viral burden on the ER can trigger the cellular unfolded protein response (UPR), which attempts to increase the protein folding and processing capacity of the ER to match the protein load. Much remains to be learned about how viruses co-opt the UPR to ensure efficient synthesis of viral glycoproteins. Here, we show that two FDA-approved thiopurine drugs, 6-TG and 6-TGo, induce the UPR in a manner that impedes viral glycoprotein accumulation for enveloped influenza viruses and coronaviruses. These drugs may impede the replication of viruses that require precise tuning of the UPR to support viral glycoprotein synthesis for the successful completion of a replication cycle.

## INTRODUCTION

Enveloped viruses encode integral membrane proteins that are synthesized and post-translationally modified in the endoplasmic reticulum (ER) prior to transport to sites of virion assembly. When ER protein folding capacity is exceeded, the accumulation of unfolded proteins in the ER causes activation of the unfolded protein response (UPR) whereby activating transcription factor-6 (ATF6), inositol requiring enzyme-1 (IRE1) and PKR-like endoplasmic reticulum kinase (PERK) sense ER stress and trigger the synthesis of basic leucine zipper (bZIP) transcription factors that initiate a transcriptional response (1). UPR gene expression causes the accumulation of proteins that attempt to restore ER proteostasis by expanding ER folding capacity and stimulating catabolic activities like ER-associated degradation (ERAD) (2). ERAD ensures that integral membrane proteins that fail to be properly folded are ubiquitinated and retrotranslocated out of the ER for degradation in the 26S proteasome. There is accumulating evidence that bursts of viral glycoprotein synthesis can burden ER protein folding machinery, and that enveloped viruses subvert the UPR to promote efficient viral replication (3, 4).

Influenza A viruses (IAVs) encode three integral membrane proteins: hemagglutinin (HA) neuraminidase (NA) and matrix protein 2 (M2). HA adopts a type I transmembrane topology in the ER, followed by addition of N-linked glycans, disulfide bond formation, and trimerization prior to transport to the Golgi and further processing by proteases and glycosyltransferases (5–11); NA adopts a type II transmembrane topology in the ER, is similarly processed by glycosyltransferases and protein disulfide isomerases, and assembles into tetramers prior to traversing the secretory pathway to the cell surface (12, 13). The small M2 protein also forms disulfide-linked tetramers in the ER, which is a prerequisite for viroporin activity (14–16). IAV replication causes selective activation of the UPR; IRE1 is activated, but PERK and ATF6 are not (17), although the precise mechanisms of regulation remain unknown. Furthermore, chemical chaperones and selective chemical inhibition of IRE1 activity inhibit IAV replication, suggesting that IRE1 has pro-viral effects. HA is sufficient to activate the UPR (18) and is subject to ERAD-mediated degradation (19). By contrast, little is known about how NA and M2 proteins affect the UPR.

Several coronaviruses (CoVs) have been shown to activate the UPR, including infectious bronchitis virus (IBV) (20, 21), mouse hepatitis virus (MHV) (22), transmissible gastroenteritis virus (TGEV) (23), human coronavirus (HCoV)-OC43 (24) and SARS-CoV-1 (25–27). However, proximal UPR sensor activation does not always elicit downstream UPR transcription responses during CoV replication (28), which suggests complex regulation of the UPR by certain CoVs. The coronavirus structural proteins Spike (S), Envelope (E) and Membrane (M) are synthesized in the ER as transmembrane glycoproteins (29). S is heavily modified by N-glycosylation in the ER, which promotes proper folding and trimerization (30). Ectopic expression of S proteins from MHV (22), HCoV-HKU1 (31) and SARS-CoV-1 (26, 31) have all been shown to be sufficient for UPR activation in cell culture. Additional transmembrane CoV proteins have been shown to be sufficient to induce the UPR, including SARS-CoV-1 non-structural protein 6 (nsp6) (32), ORF3a (33) and ORF8ab (34) proteins. CoVs rearrange ER membranes into a complex reticulovesicular network that includes double-membrane vesicles (DMVs) (35). Precise roles for these compartments remain incompletely understood, but multispanning transmembrane proteins nsp3, nsp4 and nsp6 are thought to physically anchor CoV replication/transcription complexes (RTCs) to these membranes (36, 37), and the N-glycosylation of nsp3 and nsp4 indicates the ER origin of DMVs. The observation of UPR gene products ER degradation-enhancing alpha-mannosidase-like protein 1 (EDEM1) and osteosarcoma amplified 9 (OS-9) in DMVs provides additional linkage between CoV RTCs and ER proteostasis mechanisms (38). Following replication, CoVs complete the assembly and egress process by budding into the ER-Golgi Intermediate Compartment (ERGIC) and traversing the secretory pathway. Thus, multiple stages of CoV infection are dependent on the ER.

Viral mRNAs are decoded by the host protein synthesis machinery, which could make them vulnerable to stress-induced regulation of protein synthesis. In addition to ER stress-mediated activation of PERK, three other sentinel kinases (protein kinase R (PKR); heme-regulated translation inhibitor (HRI); general control non-derepressible-2 (GCN2)) respond to diverse stresses by phosphorylating eukaryotic translation initiation factor 2-alpha (eIF2α) and enforcing a translation initiation checkpoint (39). The assembly of the ternary complex, comprised of eIF2, GTP and tRNAi^met^ is essential for incorporation of tRNAi^met^ into the 40S ribosomal subunit during translation initiation. Phosphorylated eIF2α binds to the eIF2B guanine nucleotide exchange factor and prevents it from exchanging GTP for GDP and recharging the ternary complex (40). This integrated stress response (ISR) stalls bulk protein synthesis at the initiation step and causes the accumulation of 48S preinitiation complexes and associated mRNAs that are bound by aggregation-prone RNA-binding proteins, including Ras-GAP SH3 domain-binding protein (G3BP), T-cell intracellular antigen-1 (TIA-1), and TIA-1 related protein (TIAR). These complexes nucleate cytoplasmic stress granules (SGs), sites where stalled messenger ribonucleoprotein (mRNP) complexes are triaged until stress is resolved and protein synthesis can resume (41, 42). Thus, ISR activation threatens efficient viral protein synthesis and causes the formation of antiviral SGs that impede efficient viral replication. Many viruses have acquired the means to prevent translation arrest and SG formation, and thereby limit the negative impact of ISR on viral protein synthesis.

We previously demonstrated that IAV suppresses SG formation in infected cells (43) and encodes three proteins with SG-suppressing activity; non-structural protein 1 (NS1), nucleoprotein (NP) and polymerase acidic X (PA-X) (43, 44). Treatment of IAV infected cells with protein synthesis inhibitors Pateamine A and Silvestrol caused SG formation and impeded IAV replication by inhibiting accumulation of viral proteins and downstream viral genome replication (44, 45). However, these inhibitors triggered SG formation and cytotoxic effects in uninfected cells as well, limiting their potential utility as antivirals. Because SG formation correlates with antiviral activity, we conducted an image-based high-content screen to identify molecules that selectively induce SG formation in IAV infected cells. We identified two FDA-approved thiopurine analogs, 6-thioguanine (6-TG) and 6-thioguanosine (6-TGo), that blocked IAV and HCoV-OC43 replication in a dose-dependent manner. Unlike Pateamine A and Silvestrol, these thiopurines selectively disrupted the processing and accumulation of viral glycoproteins, which correlated with UPR activation. Synthesis of viral glycoproteins could be partially restored in 6-TG treated cells by the chemical inhibition of the UPR or ISR. Our data suggest that UPR-inducing molecules could be effective host-targeted antivirals against viruses that depend on ER processes to support efficient replication. Induction of UPR by 6-TG and 6-TGo represents a novel host-directed antiviral mechanism triggered by these drugs and reveals a previously unrecognized unique mechanism of action that distinguishes them from other closely related thiopurines and nucleoside analogues.

## RESULTS

### Thiopurine analogs 6-TG and 6-TGo selectively induce SGs in IAV infected cells

To identify molecules that selectively induce antiviral SG formation in IAV infected cells without inducing SGs in uninfected cells, we developed an image-based high-content screen. Clonal A549[EGFP-G3BP1] cells (43) that stably express the core SG protein G3BP1 fused to enhanced green fluorescent protein (EGFP) were selected because they display rapid and uniform SG formation in response to known SG-inducing molecules (e.g. Silvestrol). We screened >50,000 small molecules from the Prestwick, Sigma LOPAC, and Chembridge DiverSet collections. After seeding on 96-well plates overnight, cells were infected with IAV strain A/Udorn/1972(H3N2) (IAV-Udorn) at a multiplicity of infection (MOI) of 1. At 4 hours post-infection (hpi) cells were treated with small molecules for four additional hours prior to fixation and staining with Hoechst 33342 to label nuclei. Images were automatically acquired from two channels (to detect Hoechst and EGFP-G3BP) per field and 15 fields per well of each 96-well plate using a Cellomics Array Scan VTI microscope and processed to yield values for punctate EGFP-G3BP1 intensity that reflected SG formation.

Through this screen, we identified two thiopurines, 6-thioguanine (6-TG) and 6-thioguanosine (6-TGo) (Fig. 1A), that triggered dose-dependent SG formation in IAV-infected cells (Fig. 1B). Specifically, SGs formed in approximately 10 % of 6-TG-treated or 6-TGo-treated infected cells; no SGs were detected in mock infected cells treated with either drug at the highest concentration (Fig. 1B). These findings were confirmed in parental A549 cells infected with IAV strain A/California/07/2009 (H1N1; IAV-CA/07); 6-TG treated cells displayed the formation of foci that contained SG constituent proteins G3BP1 and poly A binding protein (PABP) (Fig. 1C). These foci also contained canonical SG proteins TIAR and eIF3A (Fig. 1D), supporting their identity as *bona fide* SGs.

**Fig 1.**
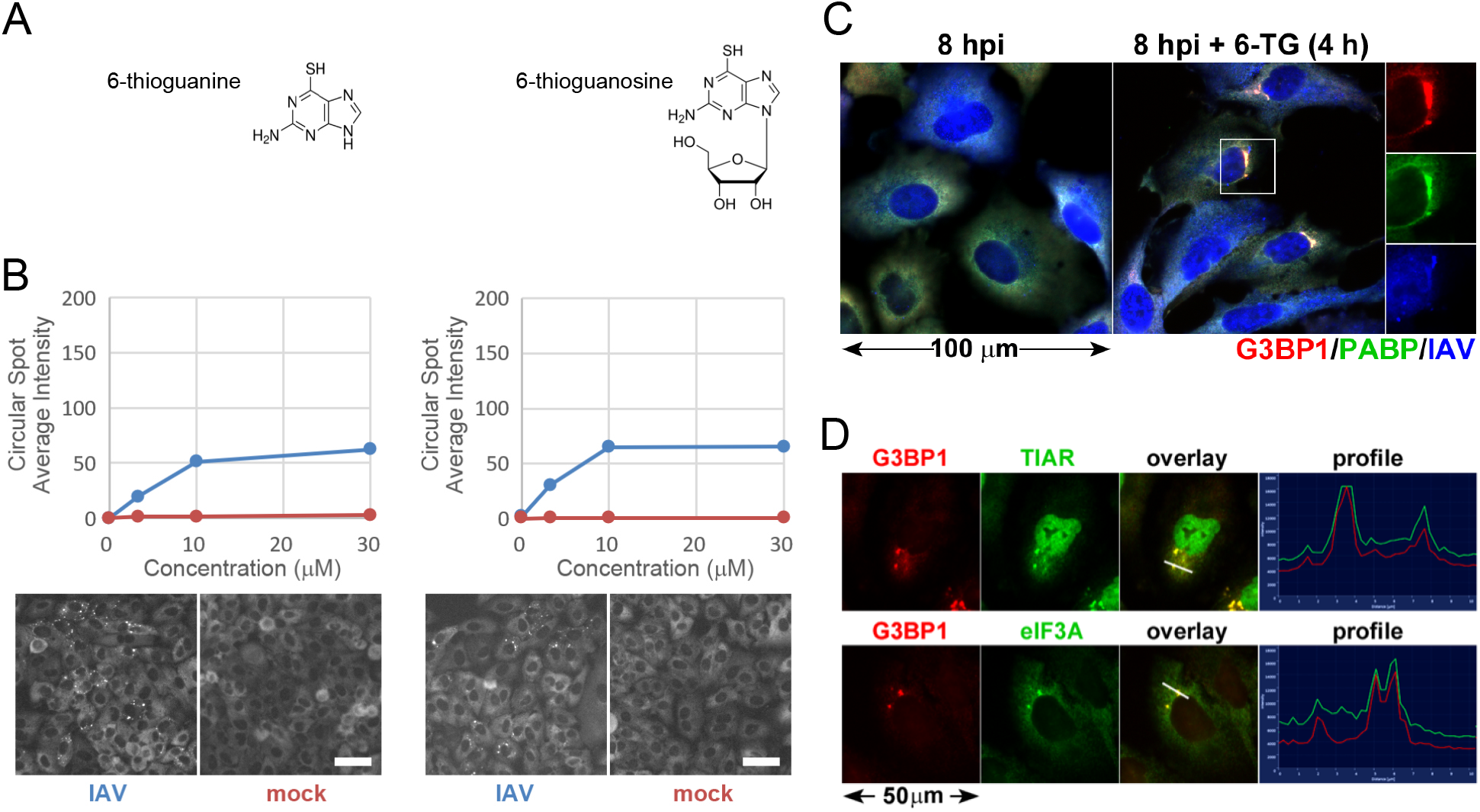
Thiopurine analogs 6-TG and 6-TGo selectively induce stress granules in IAV infected cells. (A) Structural diagrams of small molecules identified in the screen. (B) Quantification of EGFP-G3BP foci formation in IAV infected (Blue) or mock (Red) infected cells treated with increasing doses of 6-TG and 6-TGo (top) and representative Cellomics images of EGFP channel of cells treated with 30 μM 6-TG and 6-TGo (bottom). At 4 hpi, cells were treated with 0, 1, 10 and 30 uM doses of thiopurine analogs 6-thioguanine (6-TG) or 6-thioguanosine (6-TGo). At 8 hpi, cells were fixed and stained with Hoeschst 33342. Automated image capture was performed using a Cellomics Arrayscan VTI HCS reader. 15 images were captured for each well and average punctate EGFP-G3BP1 intensity was calculated. (C) A549 cells were infected with IAV-CA/07 at a MOI of 1. At 4 hpi, cells were treated with 6-TG or mock-treated. At 8 hpi, cells were fixed and immunostained with antibodies directed to stress granule marker proteins G3BP1 (red), PABP (green) and a polyclonal IAV antibody (blue) that detects antigens from NP, M1, and HA, followed by staining with Alexa-conjugated secondary antibodies. (D) A549 cells were infected with IAV-CA/07 at a MOI of 1. At 4 hpi, cells were treated with 6-TG (10μM). At 8 hpi, cells were fixed and immunostained with antibodies directed to stress granule marker proteins G3BP1 (red), TIAR (green) and eIF3A (green), followed by staining with Alexa-conjugated secondary antibodies. Images captured on a Zeiss Axioimager Z2 fluorescent microscope. Representative images shown. Scale bars represents 20 μm.

### Thiopurine analogs inhibit IAV replication

Next, we wanted to determine whether thiopurine-mediated SG formation indicated a disruption of viral replication. A549 cells were infected with IAV strain A/PuertoRico/8/1934 (H1N1; IAV-PR8) and treated with 6-TG, 6-TGo or controls at 1 hpi. Cell supernatants were harvested at 24 hpi and infectious virions enumerated by plaque assay. Despite SG induction in only a fraction of virus-infected cells, we observed a sharp dose-dependent decrease in virion production following treatment with either thiopurine analog. Treatment with 2 μM 6-TG reduced virion production by ~10-fold, whereas 2 μM 6-TGo reduced virion production by ~100-fold (Fig. 2A). Furthermore, treatment of IAV infected cells with 10 μM concentrations of either 6-TG or 6-TGo led to even greater inhibition of IAV production (Fig. 2A). This suggests that SG formation correlates with the disruption of the viral replication cycle. However, the sharp decrease in infectious virion production in 6-TG/6-TGo-treated cells suggests that SG formation is not required for their antiviral effect. The nucleoside analog 5-fluorouracil (5-FU) had no effect on IAV replication at 2 μM and 10 μM doses (Fig. 2A). Using an alamarBlue assay, we observed a ~30% reduction in A549 cell viability in the presence of 10 μM doses of 6-TG/6-TGo (Fig. 2B). Compared to SG-inducing translation inhibitor Silvestrol, which causes apoptosis in A549 cells upon prolonged exposure, we did not observe significant disruption of cell monolayer by 6-TG treatment (Fig. 2C) or induction of apoptosis as measured by PARP cleavage (Fig. 2D). This is consistent with a recent report of 6-TG-mediated cytostatic rather than cytotoxic effects on A549 cells (46). In Vero cells, 6-TG treatment partially protected cellular monolayers from IAV-induced cell death over 72-h incubation (Fig. 2E). Taken together, our data suggest that 6-TG and 6-TGo elicit a broad dose-dependent antiviral effect against IAV that was not shared by the nucleoside analog 5-FU.

**Fig 2.**
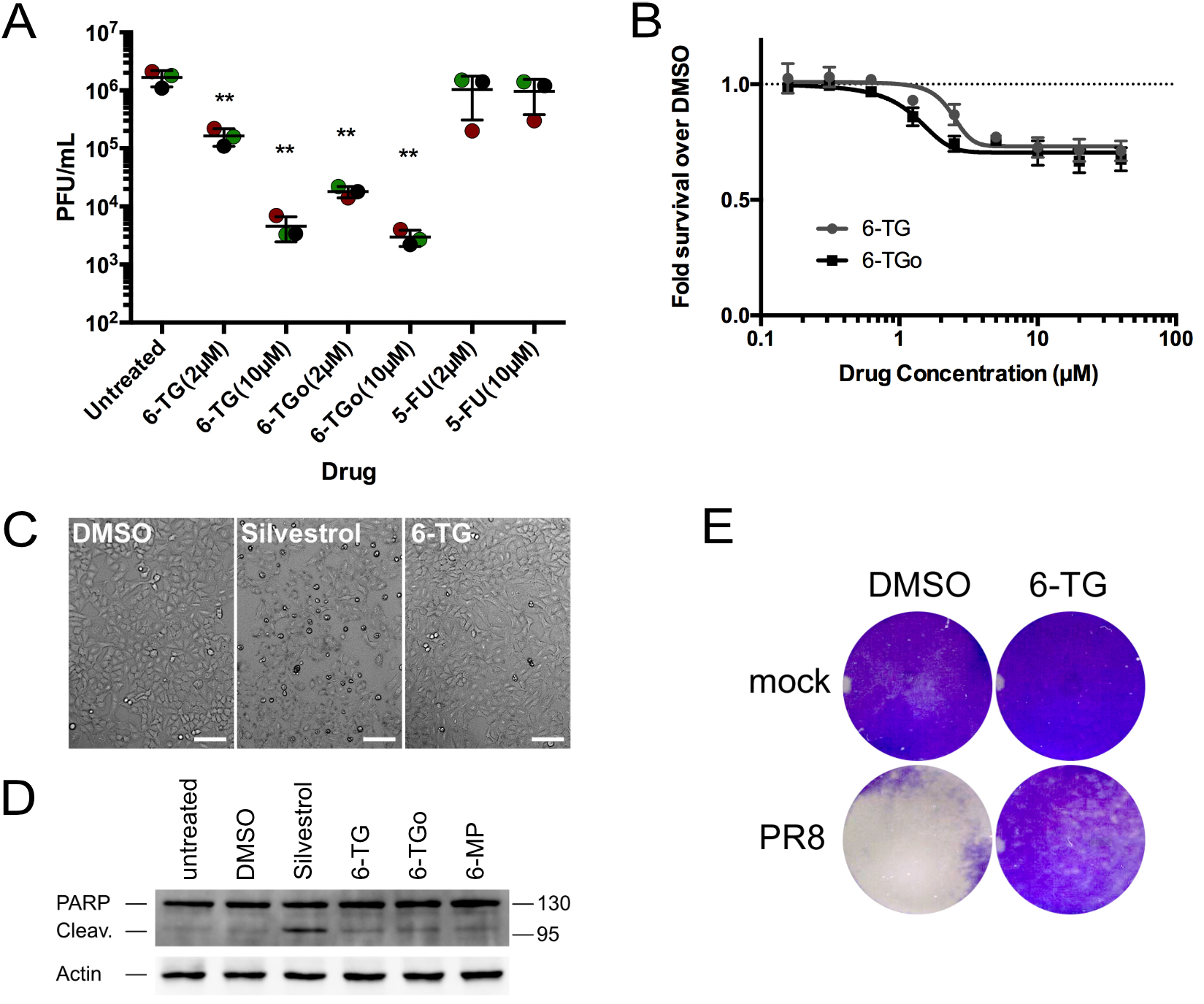
6-TG and 6-TGo inhibit IAV replication. (A) A549 cells were mock infected or infected with IAV-PR8 at a MOI of 0.1. After 1 h, cell monolayers were washed and overlaid with medium containing drugs at the indicated concentrations, or vehicle control. At 24 hpi, cell supernatants were collected and infectious virions were enumerated by plaque assay. 3 independent experiments are graphed (N=3) and error bars denote standard deviation. Circles represent biological replicates with each replicate colour coded. One-way ANOVA and post-hoc Turkey’s multiple comparisons tests were done to determine statistical significance (**, p-value < 0.01). (B) A549 cells were treated with escalating doses of 6-TG, 6-TGo, or vehicle control for 23 hours and cell viability was measured using an alamarBlue cell viability assay. Relative fluorescence units were normalized to vehicle control. Error bars represent the standard deviation (N=3). (C) Representative phase contrast images of A549 cell monolayers treated with Silvestrol (320 nM), 6-TG (10 μM), or vehicle (DMSO) control for 23 h. White scale bars in the bottom right corner of images represent 100 μm. (D) Lysates of A549 cells treated with the Silvestrol (320 nM) and thiopurines (10 μM) were analysed by western blotting for total PARP (full length and cleaved). β-actin antibody staining was used as loading control. (E) Vero cells were infected with IAV PR8 and treated with 6-TG (10μM) or vehicle (DMSO). After 72 hpi, cells were fixed with 5% formaldehyde and stained with 1% crystal violet.

### 6-TG treatment inhibits HA and NA glycoprotein processing and accumulation without affecting viral transcription

Next, we compared accumulation of viral proteins in control cells and cells treated with thiopurines and 5-FU (Fig. 3A). We previously observed that SG-inducing eIF4A inhibitors broadly affected synthesis of IAV proteins and blocked progression through the replication cycle by preventing the ‘switch’ of the RNA-dependent RNA polymerase (RdRp) from transcription to genome replication (45). By contrast, treatment of IAV-PR8 infected cells with 6-TG and 6-TGo selectively blocked the processing and accumulation of hemagglutinin (HA) and neuraminidase (NA) glycoproteins without affecting the accumulation of nucleoprotein (NP). Accumulation of matrix protein 1 (M1) was affected by higher doses of 6-TG and 6-TGo, but to a much lesser extent than HA and NA (Fig. 3A). For NA, 6-TG and 6-TGo caused a dose-dependent accumulation of faster-migrating species, reminiscent of treatment with the inhibitor of N-linked glycosylation, tunicamycin (TM). For HA, 6-TG and 6-TGo likewise caused dose-dependent accumulation of faster-migrating, presumably un-glycosylated species, but these were difficult to visualize as they migrated to the same position as NP on immunoblots probed with polyclonal anti-IAV antibodies that concurrently detect NP, M1 and HA. 5-FU, which had no effect on viral replication over the 24 h time course in these cells (Fig. 2A), likewise had no effect on the accumulation of these IAV proteins (Fig. 3A). Consistent with the notion of selective inhibition of IAV glycoprotein synthesis and maturation, we observed that 6-TG had no effect on the accumulation of IAV-PR8 *HA* or *NA* transcripts or function of the RdRp in genome replication, as 6-TG had little effect on the accumulation of HA and NA genome segments (Fig. 3B). Taken together, these data support a novel mechanism of action for thiopurine analogs in selectively inhibiting processing and accumulation of IAV glycoproteins and significantly impairing IAV replication.

**Fig 3.**
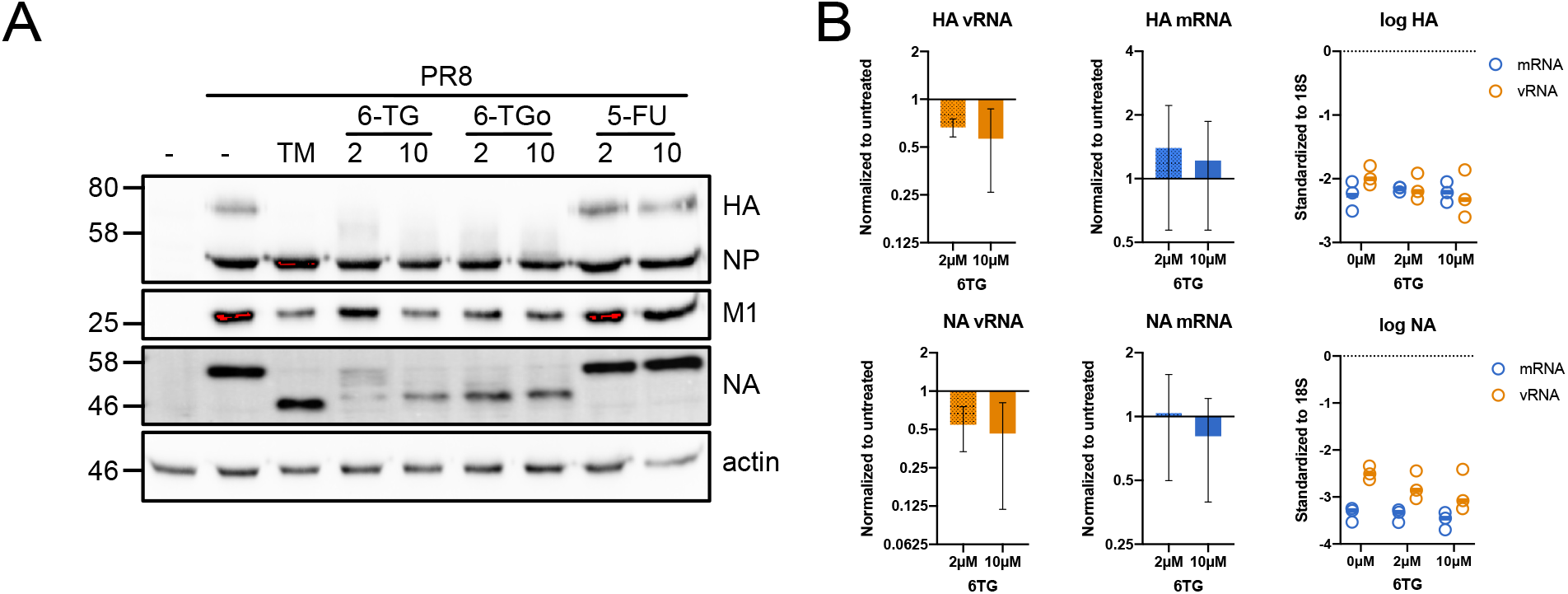
6-TG treatment inhibits HA and NA glycoprotein processing and accumulation without affecting transcript levels. (A) A549 cells were mock infected of infected with IAV-PR8 at a MOI of 0.1. After 1 h, cell monolayers were washed and overlaid with medium containing drugs at the indicated concentrations, or vehicle control. At 24 hpi, cell lysates were collected and analyzed by western blot, using a polyclonal IAV antibody that detects antigens from HA, NP and M1, or an anti-NA antibody. β-actin staining was used as a loading control. Western blots are representative of 3 independent experiments. (B) A549 cells were infected with IAV-PR8 at a MOI of 0.25, washed and overlaid with media containing 2μM or 10 μM 6-TG, or vehicle (DMSO) control. Cell lysates were collected at 24 hpi and the total RNA was isolated. The levels of IAV HA and NA mRNAs, and HA and NA genomic vRNAs, were measured by RT-qPCR. Changes in RNA levels were calculated by the ΔΔCt method and normalized using 18S rRNA as a reference gene. Error bars represent standard deviation between biological replicates (N=3); Circles represent biological replicates; Lines represent the average value.

### 6-TG and 6-TGo activate the UPR and chemical mitigation of ER stress restores synthesis of HA glycoproteins

By inhibiting N-linked glycosylation, TM impedes proper processing of secreted and transmembrane proteins in the lumen of the ER, which elicits ER stress and activates the UPR (47). Indeed, we observed that TM treatment of A549 cells caused accumulation of XBP1s and the ER chaperone binding immunoglobulin protein (BiP) (Fig. 4A). BiP upregulation is an excellent measure for UPR activation because it requires both ATF6(N)-dependent transcription of *BiP* and PERK-mediated activation of the ISR and uORF-skipping-dependent translation (1). We observed that both 6-TG and 6-TGo caused BiP and XBP1s accumulation in A549 cells, whereas the chemically similar thiopurine 6-mercaptopurine (6-MP) did not. Nucleoside analogs 5-FU and ribavirin also did not affect BiP or XBP1s levels (Fig. 4A). These data demonstrate that 6-TG and 6-TGo, but not all thiopurines, activate the UPR in A549 cells.

**Fig 4.**
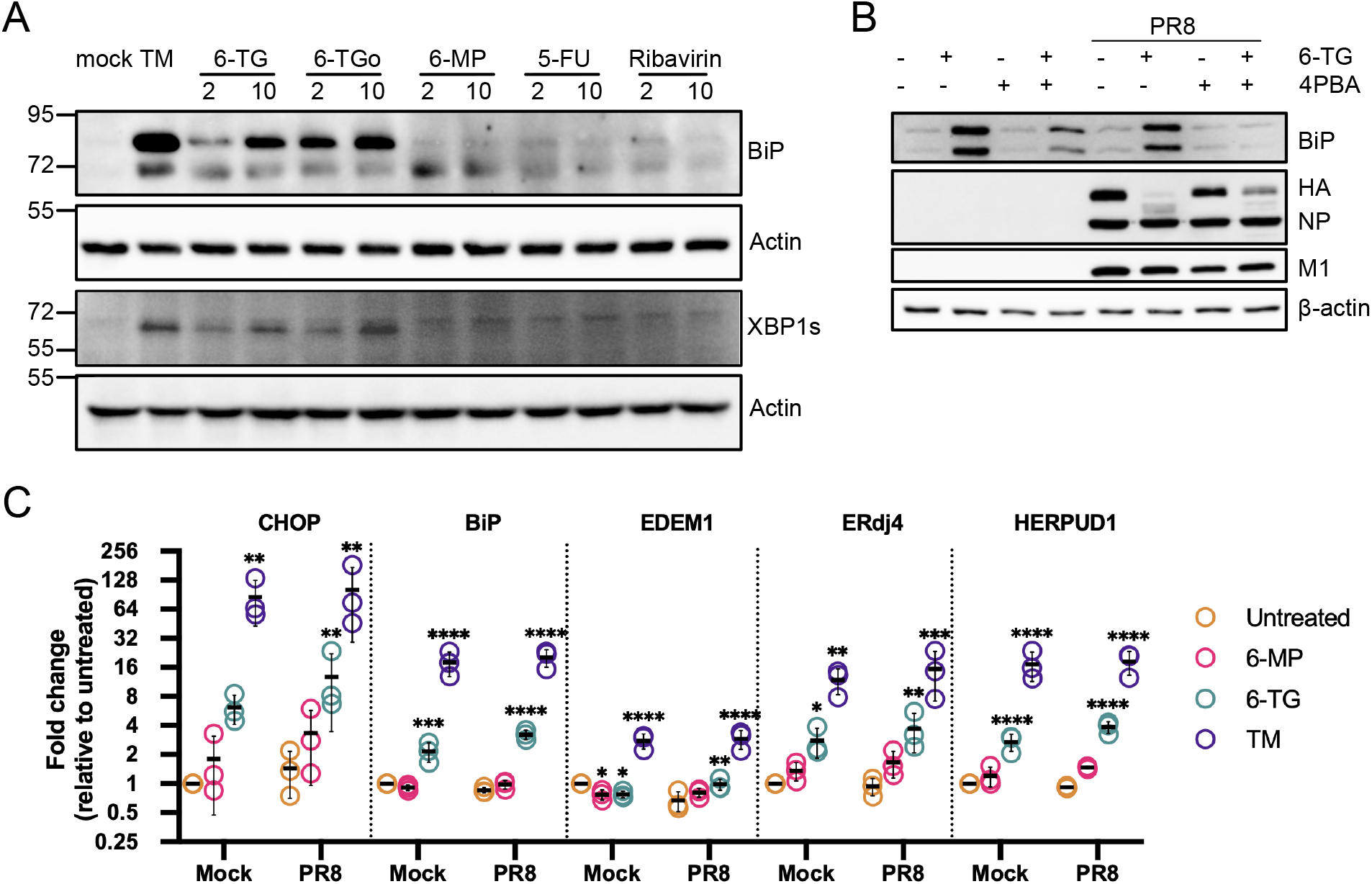
6-TG and 6-TGo activate the UPR and chemical mitigation of ER stress partially restores synthesis of HA glycoprotein. (A) A549 cells were treated with 6-thioguanine (6-TG), 6-thioguanosine (6-TGo), 6-mercaptopurine (6-MP), 5-fluorouracil (5-FU) or ribavirin at the indicated concentrations for 6 h (XBP1s) or 24 h (BiP) prior to harvesting lysates for immunoblotting. 5 μg/ml tunicamycin (TM) served as positive control for UPR activation, whereas DMSO was mock treatment. Membranes were probed with anti-BiP and anti-XBP1s antibodies to measure UPR activation. β-actin served as a loading control. (B) A549 cells were mock-infected or infected with IAV-PR8 at MOI of 1. After 1 h, cells were washed and incubated with 20 μM 6-TG or vehicle control, with or without 10 mM 4PBA, a chemical chaperone. At 20 hpi, cell lysates were harvested and probed with antibodies for the indicated target proteins. Western blots are representative of 3 independent experiments. (C) A549 cells were infected with IAV-PR8 at MOI of 1, washed and overlaid with media containing 6-MP, 6-TG, or TM. Cell lysates were collected at 24 hpi and RNA was isolated and processed for RT-qPCR. Changes in CHOP, BiP, EDEM1, ERdj4, and HERPUD1 mRNA levels were calculated by the ΔΔCt method and normalized using 18S rRNA as a reference gene and standardized to mock. Error bars represent the standard deviation between biological replicates (N=3); Circles represent biological replicates; Lines represents the average value. Statistical significance was calculated via a two-way ANOVA followed by a Dunnett multiple comparisons test.

To determine whether thiopurines could activate the UPR during IAV infection, A549 cells infected with IAV-PR8 were treated with 6-TG. We observed that 6-TG caused strong accumulation of BiP which coincided with diminished accumulation of HA (Fig. 4B). By contrast, co-administration of 6-TG and the chemical chaperone 4-phenylbutyrate (4-PBA) (48) diminished accumulation of BiP and partially restored HA levels in infected cells, without affecting levels of NP and M1 proteins. This suggests that thiopurine-mediated activation of the UPR/ISR is at least partially responsible for the diminished accumulation of HA glycoproteins in infected cells.

To corroborate our observation of 6-TG-mediated UPR activation, A549 cells were mock infected or IAV-PR8 infected, and treated with 6-TG, 6-MP, or TM for 24 h before harvesting RNA for RT-qPCR analysis of UPR gene expression. We analyzed transcripts produced from target genes linked to each arm of the UPR; ATF4 target gene *CHOP*, XBP1s target genes *EDEM1* and *ERdj4*, and ATF6(N) target genes *BiP* and *HERPUD1*. As expected, TM treatment caused strong induction of all 3 arms of the UPR and increased transcription of all five target genes in mock-infected cells and infected cells alike (Fig. 4C). This strong and consistent transcriptional output between mock-infected and infected cells suggests that the UPR remains largely intact during IAV-PR8 infection. Treatment with 6-MP had little effect on UPR gene expression (Fig. 4C). By contrast, 6-TG treatment caused statistically significant increases in transcription from all five UPR target genes (Fig. 4C). These observations confirm that 6-TG activates all three arms of the UPR, whereas the chemically similar thiopurine 6-MP does not.

### Inhibition of the integrated stress response does not restore NA processing and oligomerization in 6-TG treated cells

IAV glycoproteins are translocated into the ER, where they are modified with N-linked glycans and organize into oligomeric complexes. Upon synthesis in the ER, the type II transmembrane protein NA is glycosylated and forms dimers linked by intermolecular disulfide bonds in the stalk region (13) that then assemble into tetramers (49). We investigated the effect of 6-TG on NA processing and oligomerization using SDS-PAGE/immunoblotting procedures in the presence or absence of the disulfide bond reducing agent dithiothreitol (DTT). Since NA tetramers are known to dissociate into dimers during electrophoresis (12, 50) we annotated the ~120 kDa band as dimers/tetramers (Fig. 5A). We observed intact glycosylated NA dimers/tetramers and monomers in mock-treated IAV-PR8-infected cells, which were resolved into ~60 kDa glycosylated NA monomers in the presence of DTT (Fig. 5A). Unglycosylated NA monomers were undetectable in mock-treated cells at steady state, confirming that N-glycosylation is a rapid initial step in NA processing in the ER. TM treatment eliminated NA dimers/tetramers, leaving a minor fraction of unglycosylated NA monomers. The 6-TG treatment diminished accumulation of all forms of NA, yielding a distinct residual band that migrated closer to the size of the unglycosylated NA monomers from TM-treated cells; this suggests that 6-TG treatment interferes with proper N-glycosylation of nascent NA. Treatment with Integrated Stress Response Inhibitor (ISRIB), which prevents ISR-mediated translation arrest by maintaining eIF2B activity (51, 52), rescued accumulation of NA monomers in both TM- and 6-TG-treated cells. However, ISRIB was not able to restore NA glycosylation and oligomerization. These data provide further evidence that 6-TG inhibits IAV glycoprotein accumulation via UPR/ISR activation and extends our understanding by demonstrating that ISR suppression does not fully reverse these effects. This is further supported by our observations that administration of ISRIB alone had no impact on IAV replication while co-administration of ISRIB with TM-or 6-TG failed to restore virion production in single-cycle infection assays (Fig. 5B).

**Fig. 5.**
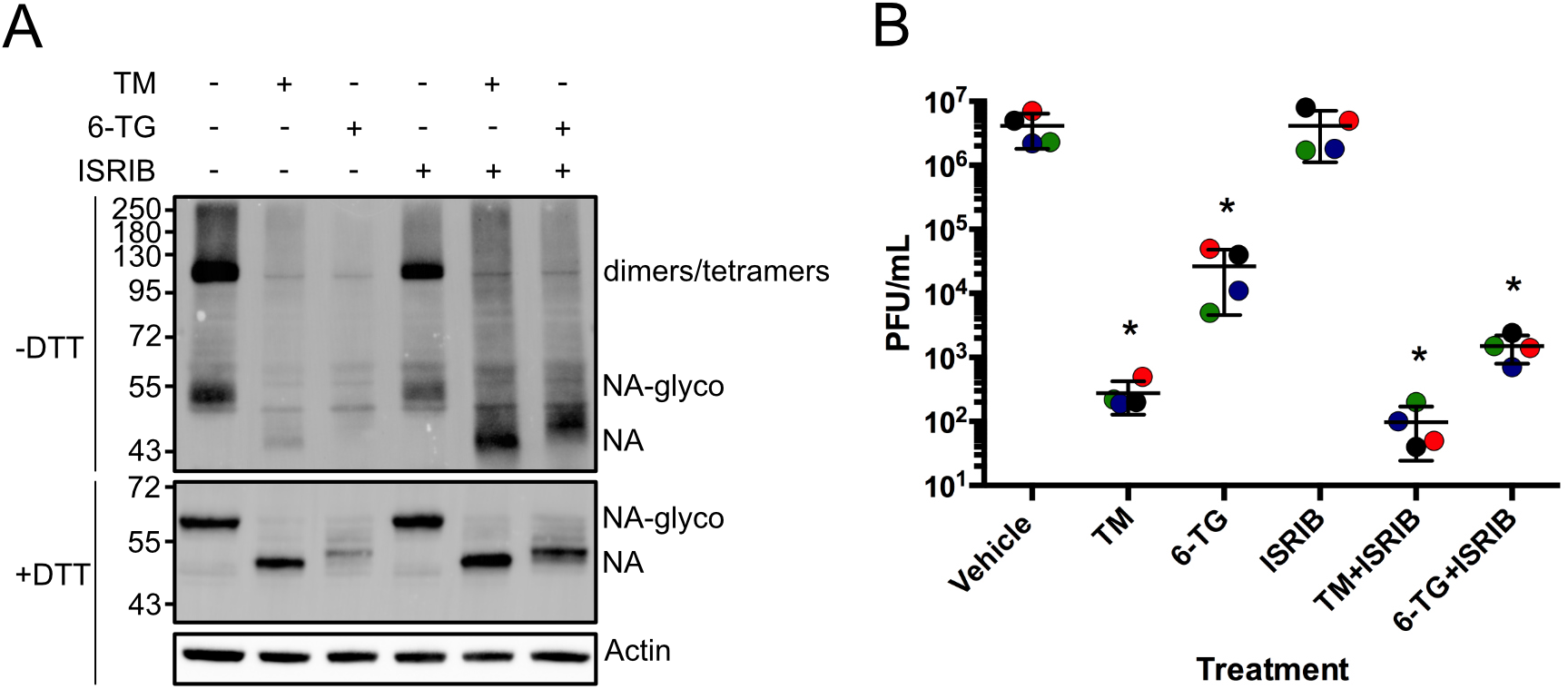
ISR inhibition restores NA synthesis in the presence of 6-TG but NA processing and virion production remain impaired. A549 cells were infected with IAV-PR8 at a MOI of 1. After 1 h, cells were washed and treated with tunicamycin (TM, 5 μg/ml), 6-thioguanine (6-TG, 10 μM) and/or 500 ng/ml ISRIB. (A) At 24 hpi, cell lysates were collected and processed for native SDS-PAGE and immunoblotting using an anti-NA antibody. N-glycosylated forms of NA are indicated as NA-glyco, whereas glycosylated NA dimers are indicated as dimers/tetramers. β-actin antibody staining was used as loading control. (B) At 24 hpi, cell supernatants were collected and infectious IAV-PR8 virions were enumerated by plaque assay. Error bars represent the standard deviation between biological replicates (N=4); Circles represent biological replicates; Lines represents the average value.

### 6-TG and 6-TGo inhibit HCoV-OC43 replication

*In vitro* studies have shown that thiopurines 6-TG and 6-MP can reversibly inhibit SARS-CoV-1 and MERS-CoV papain-like cysteine proteases PL(pro) (53–55); however, whether these thiopurines could inhibit viral replication was not assessed. Our observations of UPR activation and selective inhibition of IAV HA and NA processing and accumulation by low micromolar doses of 6-TG and 6TGo, but not 6-MP, suggest a distinct antiviral mechanism of action for these thiopurines. If true, the antiviral activity of 6-TG and 6-TGo may be broadly applicable to other viruses with envelope glycoproteins like coronaviruses. To test this directly, we performed HCoV-OC43 replication assays in human colorectal tumor-8 (HCT-8) cells in the presence of 2 μM and 10 μM doses of 6-TG, 6-TGo and 6-MP, with TM serving as a strong UPR-inducing positive control. At 24 h, cell-free and cell-associated virus was collected and combined and titered by TCID50 in Baby Hamster Kidney-21 (BHK-21) cells. We observed strong, dose-dependent inhibition of HCoV-OC43 replication in the presence of 6-TG and 6-TGo resulting in a >10-fold reduction in viral titre, whereas 6-MP had minimal effects (Fig. 6A). As expected, TM also potently inhibited HCoV-OC43 replication. Similar to A549 cells, HCT-8 cell viability was minimally affected by the thiopurines at concentrations at or below 10 μM (Fig. 6B). Consistent with the strong effect on infectious virion production, we also observed significant, dose-dependent reductions in viral protein accumulation due to 6-TG and 6-TGo treatment in HCoV-OC43-infected HCT-8 cells (Fig. 6C, the main band recognized by the anti-OC43 antibody is consistent with the size of the nucleocapsid N protein). Treatment with higher dose of 6-MP caused detectable decline in N protein levels in HCoV-OC43-infected cells, but not to the levels observed with 6-TG or 6-TGo treatment (Fig. 6C). In HCoV-OC43-infected HCT-8 cells, BiP and CHOP expression was upregulated following thiopurine treatment (Fig. 6C). This is consistent with the significant induction of BiP proten and *CHOP* mRNA levels that was observed in thiopurine-treated A549 cells (Fig. 4C). To test the effects of 6-TG on CoV replication, we analysed HCoV-OC43 mRNA synthesis by harvesting RNA from infected cells treated with 6-TG or vehicle control. We observed that 6-TG treatment caused significant decreases in steady-state levels of (+) genomic RNA (ORF1ab) as well as (+) subgenomic RNA (sgRNA) that encodes N (Fig. 6D). Thus, despite the previously reported effects of 6-TG and 6-MP on HCoV cysteine protease activity *in vitro*, we observed that 6-MP had only modest effects on HCoV-OC43 replication whereas 6-TG and 6-TGo had clear antiviral effects similar to our previous observations of inhibition of IAV replication. Thiopurine antiviral activity in our HCoV-OC43 infection assays correlated with UPR activation and hampered viral genome synthesis and viral protein production.

**Fig. 6.**
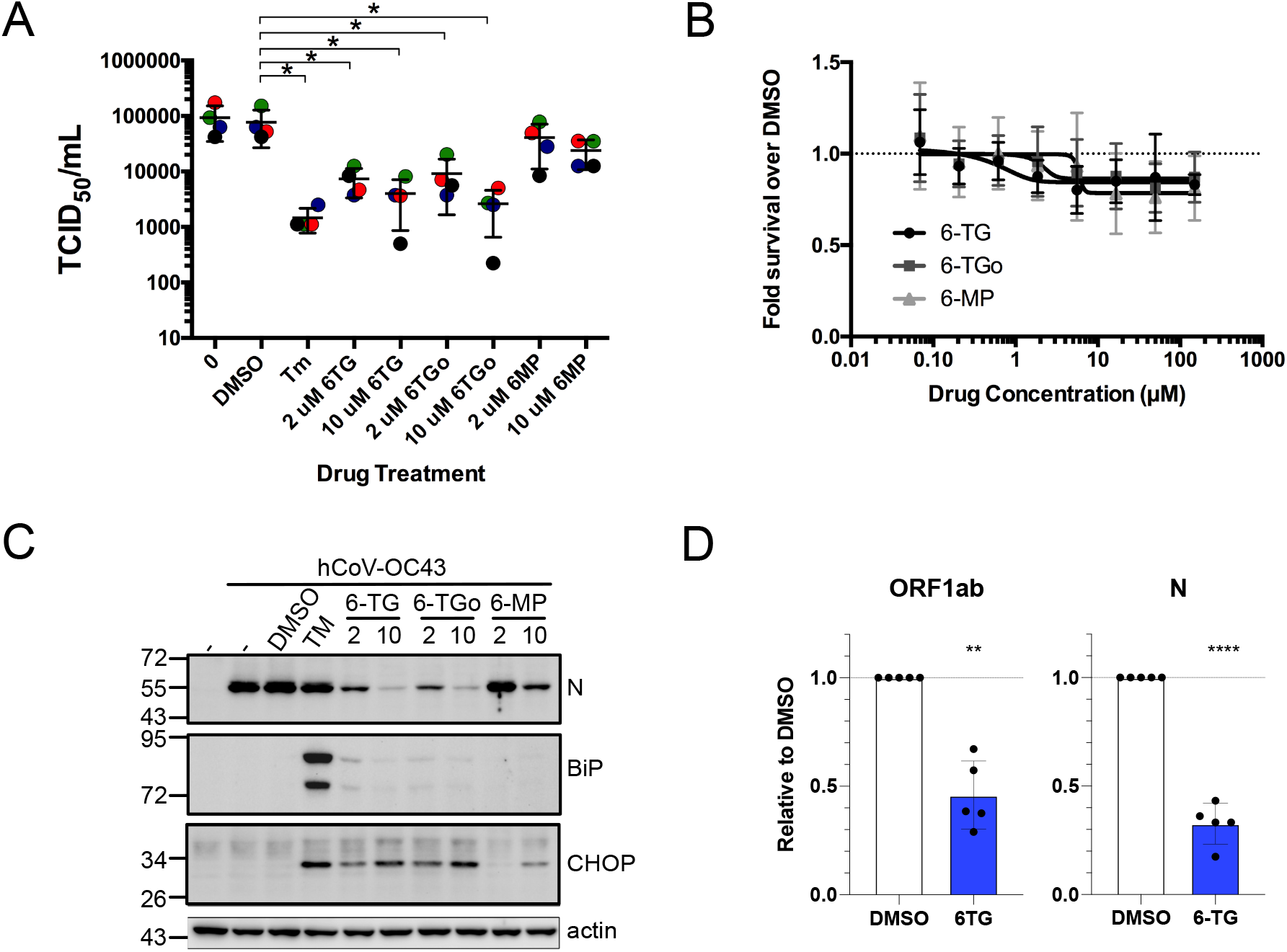
6-TG and 6-TGo inhibit HCoV-OC43 replication. Human HCT-8 cells were mock-infected or infected with HCoV-OC43 at a MOI of 0.1. At 1 hpi, cells were washed and overlaid with medium containing 2 μM or 10 μM 6-TG, 6TGo or 6-MP, or vehicle control (DMSO). (A) Progeny virus was collected at 24 hpi by combining cell-associated and secreted fractions, and titered by TCID_50_ on BHK-21 cells. Error bars represent standard deviation (N=4) and circles represent individual replicates. (B) HCT-8 cells were treated with escalating doses of the indicated thiopurines or vehicle control for 23 hours and cell viability was measured using an alamarBlue cell viability assay. Relative fluorescence units were normalized to vehicle control. Error bars represent the standard deviation (N=3). (C) Lysates of HCov-OC43 infected or mock-infected HCT-8 cells treated as indicated were analysed by western blotting. (D) RNA was harvested at 24 hpi from HCoV-OC43 infected HCT-8 cell lysates treated with DMSO or 10 μM 6-TG and processed for RT-qPCR to amplify ORF1ab and N mRNAs. Values were normalized to 18S rRNA levels (**, p-value < 0.01; ****, p-value < 0.0001, N=5).

### 6-TG inhibits SARS-CoV-2 Spike protein accumulation via UPR activation

Because 6-TG activates the UPR/ISR and inhibits the processing and accumulation of IAV glycoproteins, we reasoned that coronavirus glycoproteins would be similarly affected by 6-TG treatment. Due to the ongoing SARS-CoV-2 pandemic, numerous reagents and constructs have been rapidly developed to study this virus, including expression plasmids. We therefore sought to determine if SARS-CoV-2 S glycoprotein is sensitive to 6-TG in ectopic expression experiments. S is first translated as full-length S0 proprotein, before cleavage to S1 and S2 domains by cellular proprotein convertases like furin (56). We observed that the S protein co-localised with the ER marker calnexin when expressed alone or co-expressed with M protein (Fig. 7A). M also caused some of the S protein to accumulate in distinct regions of the cytoplasm proximal to, but not overlapping with, calnexin-stained ER, which likely represents the ERGIC (Fig. 7A). Ectopic expression of S led to accumulation of ~230 kDa full-length N-glycosylated S0 monomers; detection of ~110 kDa S1 ectodomains demonstrated efficient S N-glycosylation and trimerization in the ER and transport to the Golgi for furin cleavage (Fig. 7B). We observed that ectopic S expression was sufficient to activate the UPR/ISR, as indicated by accumulation of BiP (Fig. 7B), consistent with previous reports of SARS-CoV-1 S (26, 31). 6-TG causes a loss of membrane-bound S0 and S2, but spared the cleaved S1 subunit (Fig 7B). Co-expression of S with M altered S processing leading to different accumulation of S1 protein species, possibly due to altered S trafficking by M and retention at the ERGIC compartment (57). PNGase F treatment of the lysates to remove N-glycosylations confirmed that M and 6-TG altered glycosylation of S, but did not affect cleavage (Fig. 7C). Treatment with either the chemical chaperone 4PBA or DBeQ, a selective chemical inhibitor of the p97 AAA-ATPase, led to partial restoration of S0 and S2 (Fig 7D). Together, these observations suggest that like IAV glycoproteins, SARS-CoV-2 S glycoprotein is vulnerable to 6-TG mediated activation of the UPR/ISR and suggest a mechanism involving accelerated turnover of membrane-anchored S0 and S2 proteins by ERAD.

**Fig. 7.**
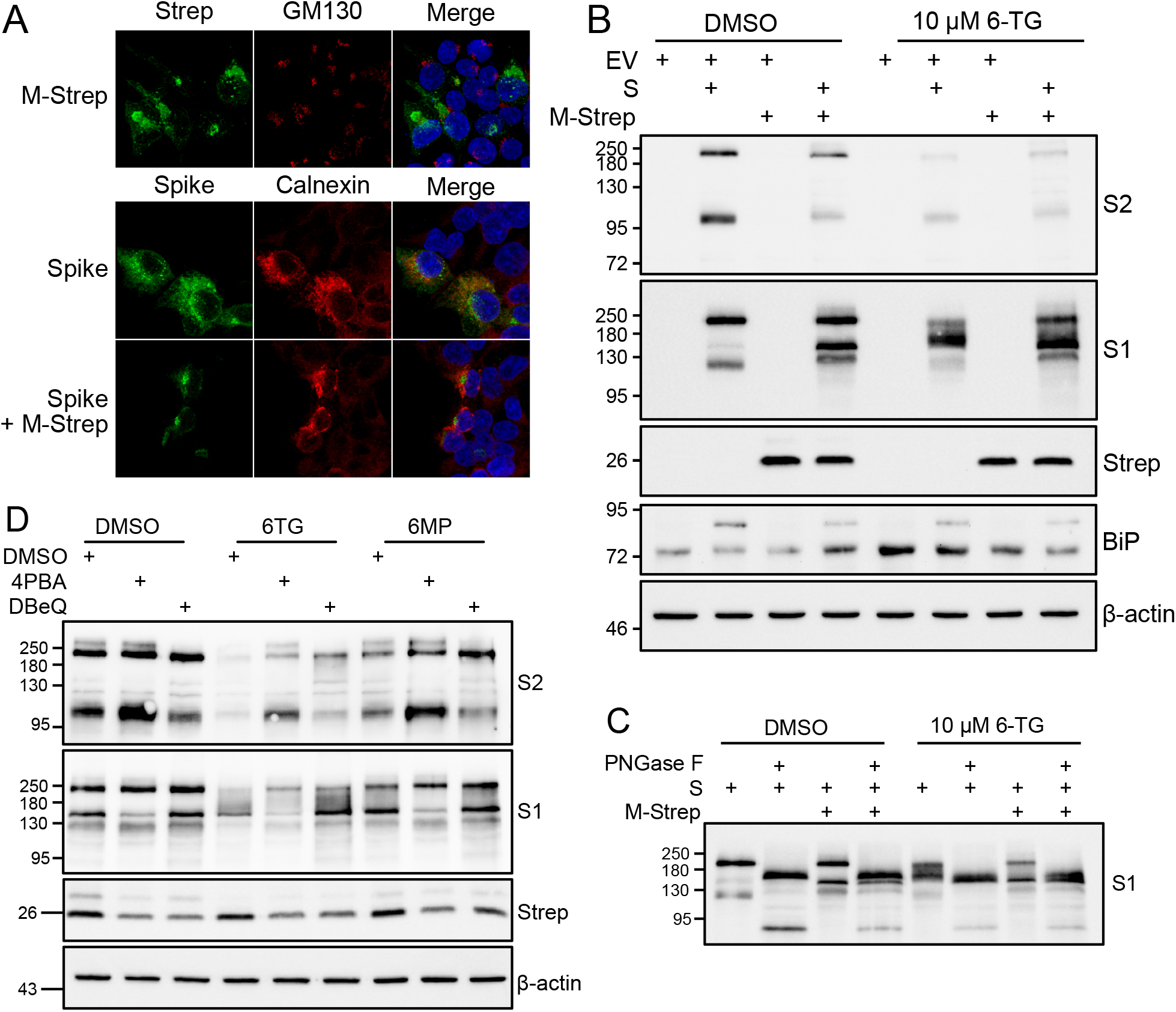
6-TG causes loss of membrane-bound SARS-CoV-2 Spike protein via ERAD. (A) 293T cells seeded on coverglass and transfected with SARS-CoV-2 Spike or 2x-strep-tagged Membrane (M-Strep) or empty vector. Cells were fixed with 4% paraformaldehyde the following day and stained as indicated. Images captured on Leica SP8, maximum intensity projections are depicted. (B) 293T cells were transfected as in (A) then treated with 10 μM of 6TG 6 h after transfection. 18 after 6TG treatment, lysates were harvested and probed by western blot as indicated. (C) Samples from A were treated with PNGase F to remove N-linked oligosaccharides. (D) 293T were transfected as in (B) and then treated with 10 μM 6TG, 10 μM 6MP, or vehicle control and co-treated 10 mM 4PBA or 10 μM DBeQ. 18h after treatment, lysates were harvested and probed by western blot as indicated.

## DISCUSSION

Compared to current direct-acting antiviral drugs, effective host-targeting antivirals may provide a higher barrier to the emergence of antiviral drug-resistant viruses. However, it remains challenging to identify cellular pathways that can be targeted to disrupt viral replication without causing adverse effects on bystander uninfected cells. Here, we report that two chemically similar FDA-approved thiopurine analogues, 6-TG and 6-TGo, have broad antiviral effects that result from activation of UPR and disruption of viral glycoprotein synthesis and maturation. Importantly, our data demonstrate for the first time that 6-TG and 6-TGo are effective antivirals against influenza virus and coronavirus and may be effective against other glycoprotein-containing viruses. 6-TG is currently used in clinical settings to treat acute lymphoblastic leukemia and other hematologic malignancies, with the main mechanism of action involving conversion into thioguanine nucleotides and subsequent incorporation into cellular DNA, which preferentially kills cycling cancer cells (58, 59). Furthermore, the active 6-TG metabolite, 6-thioguanosine 5’-triphosphate, was shown to inhibit small GTPase Rac1 (60), which is believed to be largely responsible for anti-inflammatory effects of 6-TG in treating inflammatory bowel disease (IBD) (61). Finally, 6-TG and closely related thiopurine 6-mercaptopurine (6-MP) were demonstrated to have in vitro effects against SARS-CoV-1 and MERS-CoV as reversible, slow-binding inhibitors of papain-like cysteine protease PL(pro) (53–55). Here, we demonstrate for the first time that 6-TG and 6-TGo, but not 6-MP or other nucleoside analogues (5-FU, ribavirin), induce the UPR in human cell lines. Selective disruption of viral glycoprotein accumulation in IAV-infected cells with minimal effects on other viral proteins suggests that UPR induction by 6-TG and 6-TGo is the main antiviral mechanism. Indeed, chemical chaperones and the ISR inhibitor ISRIB partially restored IAV HA, NA, and SARS-CoV-2 S protein accumulation in cells treated with 6-TG. Inhibition of the ERAD pathway with DBeQ also restored accumulation of ER membrane-anchored subunits of SARS-CoV-2 S protein (uncleaved precursor S0 and cleaved S2) in 6-TG-treated cells. This indicates that the 6-TG-induced UPR causes both the phospho-eIF2a dependent decrease in viral glycoprotein mRNA translation and the ERAD-mediated degradation of newly synthesized ER-anchored proteins. In the case of IAV which replicates in the nucleus of infected cells, depletion of viral envelope glycoproteins blocks infectious virion production but minimally affects replication of viral nucleic acids. By contrast, synthesis of coronavirus genomic RNA is inhibited by thiopurine-induced UPR. This reduction could be due to inhibition of PL(pro) activity as previously suggested (53–55); however, we suggest that thiopurines likely inhibit viral replication due to its occurrence on the multivesicular network generated from virus-rearranged ER membranes which may be highly sensitive to UPR-induced alterations.

Our analyses of IAV-infected cells did not detect induction of UPR when high levels of viral glycoproteins HA and NA accumulated at later times post-infection. Many viruses have mechanisms to interfere with different arms of the UPR, presumably to maximize the production and trafficking of viral glycoproteins, and IAV was shown to inhibit activation of PERK and ATF6 (17). Downstream, IAV blocks new host mRNA processing and protein synthesis through host shutoff mechanisms mediated by viral non-structural protein 1 (NS1) and polymerase acidic X (PA-X) proteins. Despite these viral mechanisms of suppression of host stress responses, 6-TG treatment elicited similar levels of UPR in both infected and uninfected cells, as measured by transcriptional induction of genes downstream of all three arms of UPR and accumulation of BiP protein. Thus, in our system it appears that 6-TG circumvents viral mechanisms of UPR suppression.

Despite multiple mechanisms deployed by IAV to block SG formation, 6-TG and 6-TGo treatment induced SGs in infected cells, which allowed us to identify these molecules in our image-based screen. We previously reported that in A549 cells, IAV inhibited SG formation triggered by treatment with thapsigargin, a potent inducer of ER stress, with only 9% of infected cells forming SGs compared to 35% of mock-infected cells (44). Thus, induction of SGs in approximately 10% of infected cells by 6-TG and 6-TGo treatment appears consistent with our previous observations. However, unlike thapsigargin, 6-TG and 6-TGo did not trigger SG formation in uninfected cells. The levels of UPR induction by these drugs were similar between infected and uninfected cells, highlighting that in IAV-infected cells SG formation may not be triggered exclusively by ER stress and PERK activation and may only partially contribute to antiviral effects of thiopurines. Indeed, SGs formed in a fraction of infected cells while accumulation of viral glycoproteins HA and NA was nearly completely blocked by 6-TG and 6-TGo.

Consistent with previous reports, ectopic expression of SARS-CoV-2 S protein was enhanced during M co-expression and S alone was sufficient to trigger ER stress. In this system, 6-TG treatment further potentiated UPR responses, as measured by increased BiP accumulation. Our data also highlight the sensitivity of membrane anchored viral proteins to 6-TG treatment. It is currently unknown if host glycoproteins will be similarly affected by 6-TG treatment, but this is an important question to answer due to the prevalent use of thiopurines clinically. While we suspect that 6-TG and 6-TGo will be effective against a wide-range of enveloped viruses, our future studies will investigate if SARS-CoV-2 replication can be negatively impacted following thiopurine treatment. In the midst of the current SARS-CoV-2 pandemic, repurposing FDA-approved medications as host-targeted antivirals has the potential to positively impact our treatment of coronavirus disease 2019 (COVID-19). However, further studies in cell culture models of SARS-CoV-2 infection are necessary prior to considering clinical studies.

What is the mechanism of UPR induction by 6-TG and 6-TGo? Our results suggest that the effects are unlikely to be mediated through DNA or RNA incorporation of 6-TG because 1) replicative stress does not specifically induce UPR; 2) among viral proteins, glycoprotein accumulation and processing was preferentially disrupted; 3) messenger RNA levels of HA and NA were not affected. Furthermore, the closely related thiopurine 6-MP that can be converted into 6-thioguanosine triphosphate and incorporated into nucleic acids did not induce UPR and had no effect on IAV glycoproteins or OC43 replication. Another nucleoside analogue, 5-FU, that is also incorporated into nucleic acids and can even trigger SG formation upon prolonged 48-hour incubation (62), was similarly inactive in our assays. The second previously described antiviral mechanism of action of 6-TG and 6-MP that involves direct inhibition of viral cysteine proteases is similarly unlikely to have major contribution to the observed phenotypes because UPR induction was triggered in both infected and uninfected cells and because, as mentioned above, 6-MP was not active in our assays. Thus, by process of elimination, we speculate that the mechanism of UPR induction by 6-TG and 6-TGo could involve GTPase inhibition. Numerous GTPases regulate ER homeostasis, including Rab GTPases that govern vesicular trafficking events and dynamin-like GTPases that regulate homotypic ER membrane fusion events required for the maintenance of branched tubular networks (63). Future studies will focus on identifying specific molecular targets of these UPR-inducing thiopurines using orthogonal biochemical and genetic screens.

## MATERIALS AND METHODS

### Cell lines

Human lung adenocarcinoma A549 cells, human embryonic kidney (HEK) 293T and 293A cells, human colorectal adenocarcinoma HCT-8, green monkey kidney (Vero), baby hamster kidney (BHK-21), and Madin-Darby canine kidney (MDCK) cells were all maintained in Dulbecco’s modified Eagle’s medium (DMEM; Thermo Fisher Scientific, Ottawa, ON, Canada) supplemented with 10% fetal bovine serum (FBS, Thermo Fisher Scientific, Grand Island, NY, USA) and 100 U/mL penicillin + 100 μg/mL streptomycin + 20 μg/mL-glutamine (Pen/Strep/Gln; Wisent Bioproducts, St-Bruno, QC, Canada) at 37°C in 5% CO_2_ atmosphere. HCT-8 cells were additionally supplemented with 1X MEM Non-Essential Amino Acids (Gibco). Baby hamster kidney (BHK-21) were maintained in DMEM supplemented with 5% FBS and 100 U/mL penicillin + 100 μg/mL streptomycin + 20 μg/mL-glutamine. All cell lines were purchased from the American Type Culture Collection (ATCC). Generation of A549[EGFP-G3BP1] cells is described in (43).

### Influenza viruses and infections

Viruses used in this study include A/PuertoRico/8/34/(H1N1) (IAV-PR8), A/Udorn/1972(H3N2) (IAV-Udorn), and A/California/07/2009(H1N1) (IAV-CA/07). IAV-PR8 stocks were generated using the 8-plasmid reverse genetic system, which was provided by Dr. Richard Webby (St. Jude Children’s Research Hospital, Memphis USA); IAV-Udorn was rescued from the 12-plasmid system that was provided by Dr. Yoshi Kawaoka (University of Wisconsin-Madison, Madison, USA); IAV-CA/07 was provided by the Public Health Agency of Canada (PHAC) National Microbiology Laboratory. Stocks of influenza viruses were propagated in MDCK cells in IAV infection media (DMEM supplemented with 0.5% bovine serum albumin (BSA) and 20 μg/mL-glutamine, and 1 μg/ml TPCK-treated trypsin). For infection, virus inoculums were diluted in IAV infection media without trypsin and added to cells for 1 h at 37°C. After inoculums were removed, cell monolayers were washed with PBS and fresh IAV infection media was added. Supernatants were harvested at 24 hpi unless indicated otherwise, supernatant was incubated with TPCK Trypsin at 1.5μg/mL for 1 hour at 37°C to activate influenza HA. Plaque assays were performed in MDCK cells using 1.2% Avicel (FMC, Philadelphia, PA) overlays as described in Matrosovich *et al.* (64). Cells were fixed with 5% formaldehyde at 48 hours post-infection and plaques visualised by staining with 1% crystal violet.

### Human coronavirus infections and quantitation

Stocks of human coronavirus OC43 (hCoV-OC43; ATCC, VR-1558) were propagated in Vero cells. To generate OC43 stocks, Vero cells were infected at a MOI of 0.05 for 1 h at 33°C in serum-free DMEM. After 1 h, the inoculum was removed, and the infected cells were maintained in DMEM supplemented with 1% FBS, 100 units/mL penicillin/100 μg/mL of streptomycin, and 2 mM L-glutamine (hCoV infection medium) for five days at 33°C. Upon harvest, the culture supernatant was centrifuged at 1000 x *g* for 5 min at 4°C, aliquoted, and stored at −80°C. Viral titers were enumerated by median tissue culture infectious dose (TCID_50_) in BHK-21 cells.

To test the effect of the indicated compounds on hCoV-OC43 replication, HCT-8 cells were infected with hCoV-OC43 at a MOI of 0.1 for 1 h at 33°C with the virus diluted in serum-free DMEM. After 1 h, the inoculum was removed, replaced with hCoV infection medium supplemented with DMSO or drug at the indicated concentration where indicated, and maintained at 33°C. At 24 hours post-infection, the infected cells were scraped into the culture medium and stored at −70°C. The samples were thawed on ice and mixed thoroughly by pipetting. To titre hCoV-OC43, TCID_50_ assays in BHK-21 cells were used. 50 μL of serially diluted inoculums were added to BHK-21 cells seeded into 96-well plates for 1 h at 37°C. Inoculums were removed and replaced with 200 uL of hCoV infection medium and incubated for five days at 37°C. After five days, the culture medium was removed, the cells washed once with PBS, fixed with 100% methanol for 15 minutes at room temperature, followed by staining with 1% crystal violet. The TCID_50_ assay were imaged and the viral titres determined using Spearman-Karber method.

### Cellomics drug screen

Generation of A549[EGFP-G3BP1] cells stably expressing EGFP-tagged G3BP1 is described in (43). A549[EGFP-G3BP1] cells were seeded at 50,000 cells/well in 96 well optical plates in the 50 μL of DMEM containing 10% FBS at 18 h prior to infection. Cells were infected with IAV-Udorn (H3N2) at a multiplicity of infection (MOI) of 1.0 by direct addition of 50 μL/well of virus inoculum pre-diluted in DMEM containing 0.5% BSA. Cells were treated with a small molecule library (~50,000 molecules) at 4 h post infection (hpi) using a pinning robot. At 8 hpi (4 h post drug treatment) cells were washed with PBS before fixation with 3% PFA in PBS supplemented with 500 ng/ml Hoechst 33342 for 20 minutes, followed by one more PBS wash. Automated image capture was performed using a Cellomics Arrayscan VTI HCS reader. Punctate EGFP-G3BP1 pixel intensity was acquired as ‘circ spot average intensity’ using the instrument’s Thermo Scientific Compartmental Analysis bioapplication. Every plate contained uninfected, untreated cells for negative control and cells treated with Silvestrol (300 nM) for positive control of SG induction. Candidate hits that were initially identified in the screen were followed up by treating both mock and IAV-Udorn infected A549[EGFP-G3BP1] cells and analyzing G3BP1 puncta.

### Chemical Inhibitors

6-thioguanine (6-TG), 2-Amino-6-mercaptopurine riboside hydrate (6-TGo), 6-mercaptopurine (6-MP), 5-fluorouracil (5-FU), Ribavirin, 4-Phenylbutyric acid (4-PBA), Integrated stress response inhibitor (ISRIB), DBeQ,, and Tunicamycin (TM) (all obtained from Sigma-Aldrich Canada Co., Oakville, ON, Canada) were solubilized in dimethyl sulfoxide (DMSO) and stored at −70°C. Stock concentrations were diluted to the indicated concentrations in media.

### Cytotoxity assay

A549 or HCT-8 cells were seeded at 10,000 cells/well in a 96 well plate. Drugs were diluted at indicated concentrates and incubated with the cells for 24 hours. At 20 hours post treatment, 10% alamarBlue Cell Viability Reagent (ThermoFisher, DAL1025) was added and further incubated for 4 hours. Plates were read on FLUOstar Omega 96 well plate reader at an excitation of 544 nm, and an emission of 580-590 nm. Both Vehicle (DMSO) and drug treatments were normalized to untreated cells. Drug treatments were then normalized to DMSO.

### Immunoblotting

Cell monolayers were washed once with ice-cold PBS and lysed in 2x Laemmli buffer (4% [wt/vol] sodium dodecyl sulfate [SDS], 20% [vol/vol] glycerol, 120 mM Tris-HCl [pH 6.8]). DNA was sheared by repeated pipetting with a 21-gauge needle before 100 mM dithiothreitol (DTT) addition and boiling at 95°C for 5 min. Samples were stored at −20°C until analysis. Total protein concentration was determined by DC protein assay (Bio-Rad) and equal quantities were loaded in each SDS-PAGE gel. For glycosylation analysis, lysates were mixed with 10X GlycoBuffer 2, NP-40, and PNGaseF (Cell Signaling Technolgies), and incubated at 37°C for 1 h before analysis by SDS page. Proteins were transferred to polyvinylidene difluoride (PVDF) membranes (Bio-Rad) with the Trans-Blot Turbo transfer apparatus (Bio-Rad). Membranes were blocked with 5% bovine serum albumin or 5% skin milk in TBS-T (Tris-buffered saline, 0.1% [vol/vol] Tween) before probing overnight at 4°C with antibodies raised to the following targets: goat polyclonal antibody to influenza A virus (ab20841, Abcam Inc., Toronto, ON, Canada), Mouse anti influenza A NA 1 (GeneTex, GT288), Mouse anti XBP1s (Cell signaling, 12782), Mouse anti CHOP (Cell signlaing, 2895), Rabbit anti SARS COV-2 S1 RBD (Elabscience, E-AB-V1006), Rabbit anti PARP (Cell signaling, 9542), Rabbit anti BiP (Cell signaling, 3177), and β-actin (13E5, HRP-conjugated, NEB). Membranes were washed with TBS-T and incubated with HRP-linked secondary antibodies prior to detection with Clarity-ECL chemiluminescence reagent (Bio-Rad). All blots were imaged on a Bio-Rad ChemiDoc-Touch system. Molecular weights were determined using protein standards (New England Biolabs P7719).

### SARS CoV-2 protein over expression

HEK 293T cells were transfected with pcDNA3.1 (EV), pLVX-M-2xStrep [Gordon et al..,2020], or pcDNA3.1 with a codon-optimized S (GenScript) using polyethylenimine. Cells were transfected in serum free DMEM for 6 h before media was replaced with 10% FBS DMEM and treated with DMSO or chemical inhibitors as indicated.

### Immunofluorescence microscopy

For immunofluorescence microscopy, A549 cells were seeded on glass coverslips and cultured overnight prior to IAV-CA/07 infection at MOI of 1, or mock infection. At the indicated times post-infection, cells were fixed with 4% paraformaldehyde and permeabilized with cold methanol as described in (65). HEK293T cells were grown on poly-D-lysine coated coverslips and transfected as described above. Cells were stained with goat polyclonal antibody to influenza A virus (ab20841, Abcam Inc., Toronto, ON, Canada, mouse-anti G3BP1 (BD Biosciences, 611126), Mouse anti-PABP (Santa Cruz, 10E10), Rabbit anti-TIAR (Cell signaling, 8509), 1:3000 rabbit anti-GM130, 1:100 rabit anti-Calnexin, 1:1000 anti-Spike (GeneTex), 1:100 anti-StrepTag (IBA), and 1:800 anti-eIF3A (Cell signaling, 3411), followed by Alexa-coupled donkey or goat secondary antibodies (Thermo Fisher Scientific) at 1:1000 dilution. Nuclei were stained with Hoechst 33342 dye. Images were captured using Zeiss AxioImager Z2 microscope or an Leica SP8 confocal microscope.

### Viral Gene Expression and Genome Replication

RNA was extracted from infected cells using the RNeasy Plus Mini Kit (Qiagen Inc., Toronto, ON, Canada) and cDNA was generated using Maxima H Minus Reverse Transcriptase (Thermo Fisher Scientific, Grand Island, NY, USA) in separate reactions containing the gene-specific primer for 18S rRNA (5’-AGGGCCTCACTAAACCATCC-3’) and either the influenza A virus-specific universal primer Uni12 (5’-AGCAAAAGCAGG-3’, for vRNA) or the oligo(dT)18 primer (for mRNA and HCoV-OC43 genomic RNA). Segment specific primers were used to analyze IAV genomic RNA and mRNA. Quantitative PCR analysis was performed using PerfeCta SYBR Greeen Fast Mix (Quanta Bio). Relative initial template quantities were determined using the ΔΔCt method. Amplifications were performed using Bio-Rad CFX Connect instrument and analyzed using the Bio-Rad CFX Manager 3.1 software. Primer sequences are presented in Table 1.

**Table 1.**
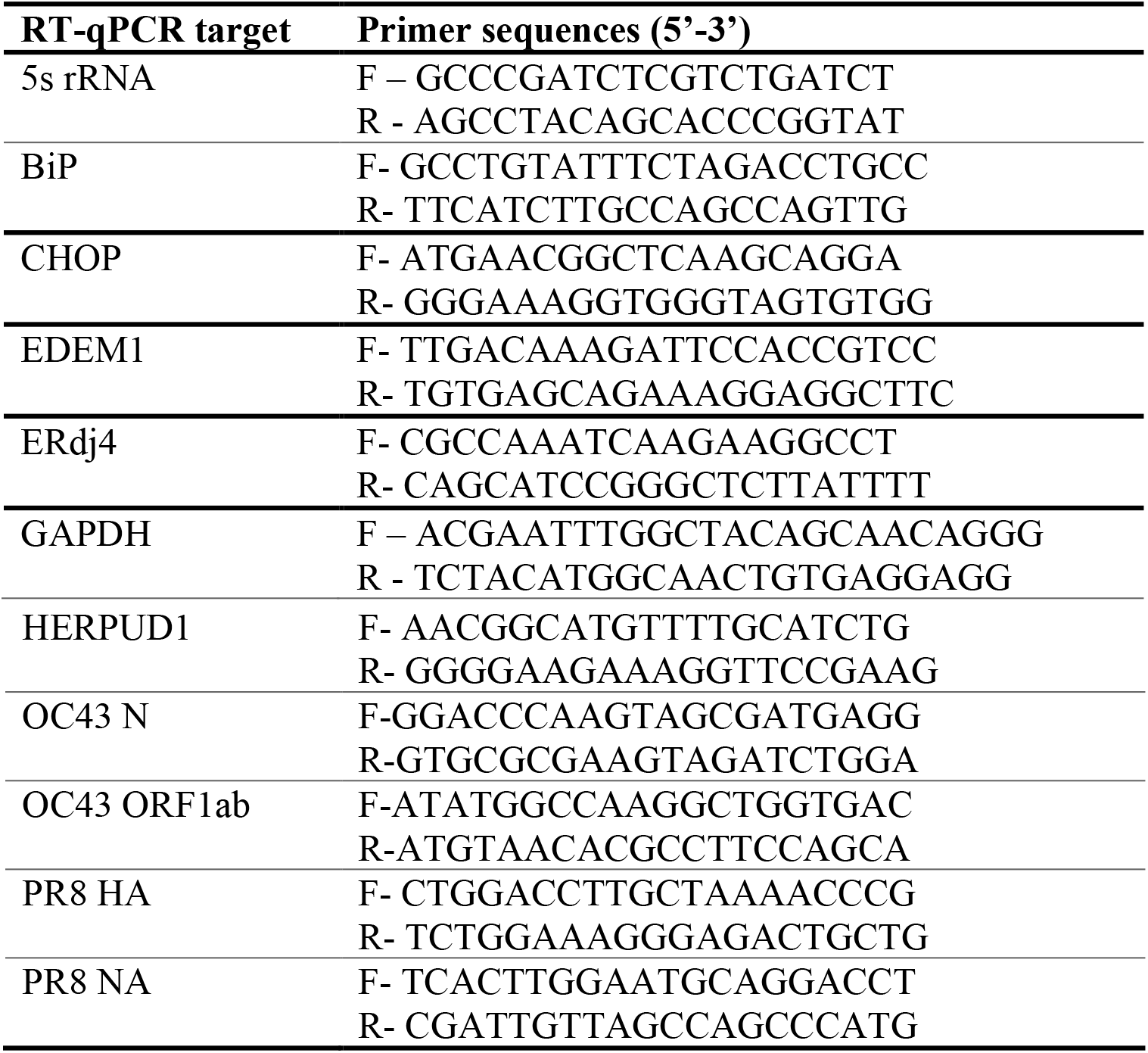
Primer sequences for RT-qPCR analysis

### Statistical analysis

Statistical analysis was performed using PRISM graph pad 8, using a one-way ANOVA multiple comparison test (Tukey) or Two-way ANOVA using the multiple comparison test using Dunnett’s correction. Significance is indicated with * (p-value of <0.05), ** (p-value of <0.01), *** (p-value of <0.001), **** (p-value of <0.0001).

## ACKNOWLEDGEMENTS

We thank members of the McCormick and Khaperskyy labs for critical reading of the manuscript. We thank Richard Webby (St. Jude Children’s Research Hospital), Yoshihiro Kawaoka (University of Wisconsin-Madison) and Todd Hatchette (Dalhousie University) for reagents. This work was supported by Canadian Institutes for Health Research grants MOP-136817 (to C.M.).

